# GLMMcosinor: Flexible Cosinor Modeling to Characterize Rhythmic Time Series Using a Generalized Linear Mixed Modeling Framework

**DOI:** 10.1101/2024.04.10.588934

**Authors:** Rex Parsons, Oliver Jayasinghe, Nicole White, Prasad Chunduri, Oliver Rawashdeh

**Affiliations:** Australian Centre for Health Services Innovation and Centre for Healthcare Transformation, School of Public Health and Social Work, Faculty of Health, Queensland University of Technology, Kelvin Grove, Australia; School of Biomedical Sciences, Faculty of Medicine, The University of Queensland, Brisbane, St Lucia, Australia

**Keywords:** Rhythmic data analysis, Cosinor modeling, Circadian rhythms, Biological rhythms

## Abstract

**Background:** Modeling rhythmic biological processes, such as gene expression and sleep-wake cycles, is critical for understanding physiological mechanisms and their dysregulation in disease. Traditional cosinor analysis, commonly used to model rhythmic data, assumes Gaussian-distributed residuals and does not account for hierarchical data, limiting its applicability in modern biological datasets.

**Results:** We present GLMMcosinor, an R package that integrates cosinor modeling into the Generalized Linear Mixed Modeling (GLMM) framework using glmmTMB. GLMMcosinor enables analysis of a broad spectrum of non-Gaussian and hierarchical data structures, including count, positive-only, and zero-inflated distributions. By incorporating mixed-effects modeling, GLMMcosinor improves parameter estimation and biological interpretability. The package includes functions for group comparisons of rhythmic parameters and visualization tools such as polar and time series plots. Additionally, GLMMcosinor is available as a Shiny app for intuitive, code-free analysis.

**Conclusions:** GLMMcosinor significantly extends the flexibility and scope of rhythmic data analysis by incorporating GLMM functionality. It is freely available on GitHub, CRAN, rOpenSci, and the R-universe, with comprehensive documentation and reproducible examples, making it a robust tool for researchers analyzing complex rhythmic datasets.

## Background

Biological rhythms underlie key physiological functions, including sleep, metabolism, and hormonal regulation. Circadian rhythms (∼24-hour cycles) are generated by endogenous oscillators synchronized to environmental cues, such as light and feeding schedules(1, 2). These rhythms are orchestrated by the central circadian clock, enabling organisms to anticipate and adapt to environmental fluctuations. Infradian and ultradian rhythms operate on longer and shorter cycles, respectively(3, 4).

Accurate quantification of rhythmic parameters—such as amplitude, period, and acrophase— is essential for understanding biological mechanisms and their variability under different conditions. For example, disruptions in circadian rhythms have been implicated in numerous diseases, including metabolic disorders, cancer, and neuropsychiatric conditions(5-7).

Cosinor analysis is a well-established method for modeling periodic data, fitting a sinusoidal function to estimate the midline statistic of rhythm (MESOR), amplitude, and acrophase(8, 9). Despite its widespread use, standard cosinor analysis has two key limitations. Traditional cosinor models assume Gaussian residuals and don’t allow for nested data structures, which may not always be appropriate for biological datasets, particularly those from repeated measures experiments or those with count or positive-only response variables(10-12).

GLMMcosinor addresses these limitations by employing a Generalized Linear Mixed Model (GLMM) framework, which supports a broader range of response distributions and hierarchical data structures. The package accommodates exponential family distributions, including Gaussian, Poisson, and Gamma, making it highly adaptable to diverse biological and medical datasets.

### Design and Implementation

GLMMcosinor extends traditional cosinor modeling(9, 13) by leveraging the glmmTMB package for fitting both Gaussian and non-Gaussian data. This flexibility allows researchers to analyze a wide range of biological datasets, including continuous, positive-only, or count data. For instance, gene expression data, which are constrained to positive values, may be better modeled using Gamma or log-normal distributions rather than a Gaussian distribution. Similarly, discrete biological data, such as cell counts or neural spike events, often follow Poisson or Negative Binomial distributions. By incorporating these additional distributional options, GLMMcosinor provides a better representation of the data generating process, which enhances biological interpretability in fields such as chronobiology, endocrinology, and physiology.

Many biological datasets contain a high proportion of zero values, which traditional cosinor models cannot effectively handle. GLMMcosinor can fit zero-inflated models, which better handles these zeros counts, improving the representation of sparse datasets with frequent zeros—particularly in behavioral and cellular biology, where infrequent actions or episodic events may dominate.

Hierarchical structures are common in biological research, often involving repeated measures on the same subjects, comparisons across tissues, or longitudinal observations. Mixed-effects models separately estimate both fixed effects (e.g., rhythmic characteristics and treatment effects) and random effects (e.g., subject- or tissue-specific variability), thereby improving the robustness of statistical inferences.

One core objective in rhythmic studies is comparing rhythmic parameters across experimental conditions—such as assessing how glucose rhythms vary with feeding schedules or evaluating pharmacological effects on circadian cycles. GLMMcosinor provides dedicated functions for testing group differences in amplitude, phase, and other rhythmic parameters. This functionality facilitates valuable insights into biological and clinical implications of temporal variations, including biomarker identification and treatment optimization(14).

### Model Specification and Interpretation

GLMMcosinor allows users to specify cosinor models using a formula-based syntax, similar to lme4(15).

1. Nonlinear form:

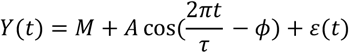
2. Linearized form:

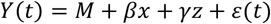

Where:

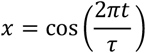

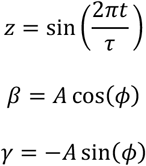

The nonlinear cosinor model (1) is linearized to allow fitting within a GLMM framework (2). In this model specification, the parameters β and γ can be used to recover the rhythm’s amplitude and phase using the delta method(16).

GLMMcosinor also supports hierarchical modeling, including random effects for within-subject correlations associated with repeated measures. To demonstrate GLMMcosinor’s utility, we include an example analyses section below where we include two sets of analyses. Firstly, we reanalyzed rhythmic gene expression data previously modeled under Gaussian assumptions. By fitting a Gamma distribution, we restricted predictions to positive values, yielding biologically realistic estimates. Secondly, to demonstrate the advantage of using a mixed-model where appropriate, we also include an example analysis comparing these two modelling approaches with an experimental dataset with repeated measures. These analyses underscore the importance of selecting appropriate distributions in cosinor modeling, especially for non-Gaussian biological data, using mixed-models when fitting models to hierarchical data. For both example analyses, we compared model fit using relevant visualizations and three key metrics: the Akaike Information Criterion (AIC), Bayesian Information Criterion (BIC), and the Residual Mean Squared Error (RMSE).

### Comparative Evaluation with Existing Tools

Despite the wide adoption of cosinor-based methods in rhythmic data analysis, most existing tools are constrained by restrictive modeling assumptions. These include reliance on Gaussian residuals, limited support for hierarchical (nested or repeated-measures) data structures, and a lack of flexibility in modeling diverse data types commonly found in biological research. To highlight the methodological advancements introduced by GLMMcosinor, we compared it against several widely used rhythm modeling tools: CircaCompare(17), Cosinor, DiscoRhythm(18), FMM, Kronos, LimoRhyde(19), and RhythmCount(20).

Each of these tools provides a subset of functionality for modeling rhythmic phenomena, yet most fall short when applied to complex biological datasets.

In Table 1, we provide a structured overview comparing the core strengths, limitations, and the added value provided by GLMMcosinor across these tools. Key dimensions of comparison include support for generalized linear models (GLMs), hierarchical modeling, compatibility with non-Gaussian distributions (e.g., Gamma, Poisson), multi-component rhythmic modeling, and inclusion of covariates within the model specification.

**Table 1.** Comparison of GLMMcosinor with existing rhythmic modeling tools across key methodological features. This table summarizes the capabilities of GLMMcosinor relative to seven widely used tools for analyzing biological rhythms: CircaCompare, Cosinor, DiscoRhythm, FMM, Kronos, LimoRhyde, and RhythmCount. Tools are evaluated based on four criteria: support for generalized linear mixed modelling/non-gaussian data, hierarchical modeling, multi-component rhythmic modeling, and inclusion of covariates in the model structure. A checkmark (✓) indicates support for the feature. GLMMcosinor is the only tool supporting all four capabilities, highlighting its flexibility and suitability for modeling complex biological time-series data. ^*^ indicate a limitation: restricted to Poisson/NB.

| Feature | GLMMcosinor | CircaCompare | Cosinor | DiscoRhythm | FMM | Kronos | LimoRhyde | RhythmCount |
| --- | --- | --- | --- | --- | --- | --- | --- | --- |
| GLM framework/ Non-Gaussian Data | ✓ | ✗ | ✗ | ✗ | ✓ | ✗ | ✗ | ✓* |
| Hierarchical modeling | ✓ | ✓ | ✗ | ✗ | ✗ | ✗ | ✗ | ✗ |
| Multi-component rhythms | ✓ | ✗ | ✗ | ✗ | ✓ | ✓ | ✓ | ✓ |
| Additional covariates | ✓ | ✗ | ✗ | ✗ | ✗ | ✓ | ✓ | ✗ |

Together, these comparisons emphasize that GLMMcosinor fills a critical gap in the rhythm modeling landscape. Its ability to accommodate hierarchical data structures, model non-Gaussian distributions, and flexibly incorporate covariates makes it uniquely suited for the complexities of modern biomedical data. The combination of statistical rigor, modeling breadth, and user accessibility establishes GLMMcosinor as a tool for analyzing biological rhythms across a wide range of disciplines.

## Results

To demonstrate the importance of selecting an appropriate modeling approach for rhythmic datasets, we compared the analysis of previously reported Liver *Nr1d1* gene expression data(21) using two models: a standard cosinor model and a GLMMcosinor model with a more suitable Generalized Linear Model (GLM) family for the response variable. The original study analyzed gene expression rhythms in two experimental groups of mice: daytime restricted feeding (DRF) and nighttime restricted feeding (NRF)(21).

A key issue with the standard cosinor model was its assumption of a Gaussian residuals, which led to biologically implausible estimates—specifically, gene expression levels below zero for the NRF group. To demonstrate this, we refit the model using GLMMcosinor under both Gaussian and Gamma distributions (Figure 1), ensuring that the response variable remained strictly positive in the Gamma model. Both models were fit using restricted maximum likelihood, and the Gamma model yielded superior performance, improving all three model fit criteria (AIC, BIC, and RMSE) (Table 2).

**Table 2.**
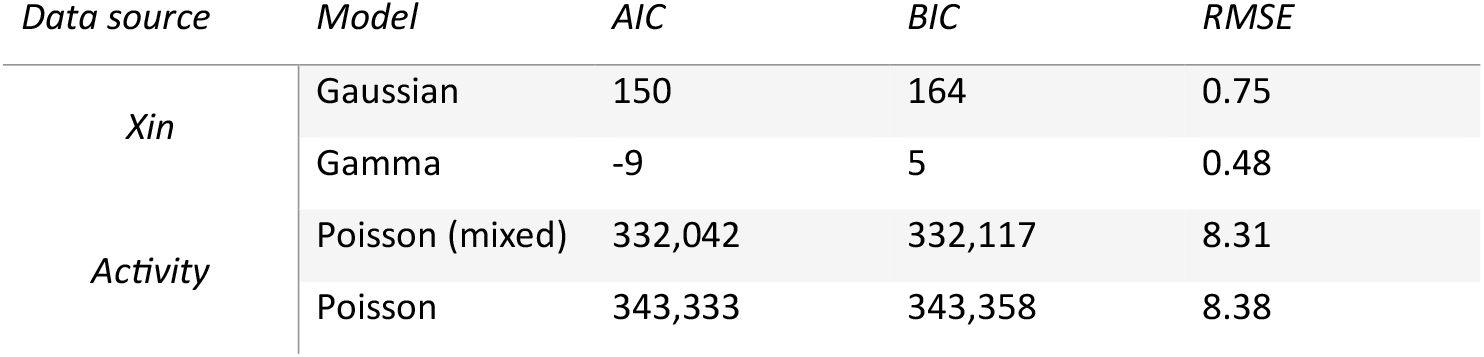
Model metrics for example analyses.

**Figure 1.**
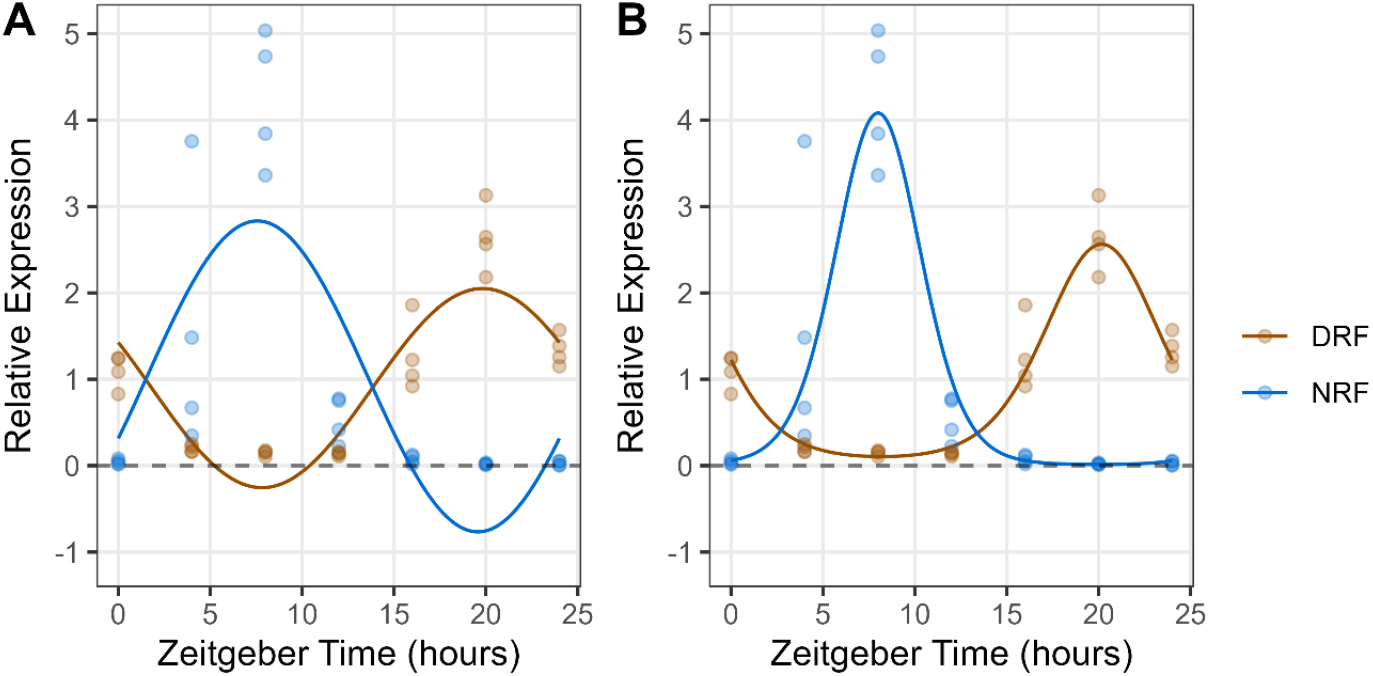
Nr1d1 relative gene expression for mice under daytime restricted feeding (DRF) and nighttime restricted feeding (NRF) conditions. Cosinor models shown in each plot were fit using Gaussian (A) and Gamma (B) families.

To our knowledge, no other R or Python package currently supports cosinor modeling with a Gamma distribution. At the time of the original publication, this issue would have arisen with any available software, underscoring the unique advantage of GLMMcosinor in handling these types of rhythmic data.

### Appropriate use of mixed models

To illustrate the importance of incorporating random effects in cosinor modeling, we analyzed activity data from 10 mice, comparing models with and without adjustments for repeated measurements from each mouse. In these data, each mouse serves as a grouping structure, and observations are not independent and identically distributed due to intra-individual correlation. In this case, a mixed-effects model is appropriate.

We used a Poisson family and compared a mixed-effects Poisson cosinor model to a naïve (non-mixed) Poisson cosinor model. Both models were fit using restricted maximum likelihood.

Figure 2 shows the time-series for each mouse in a separate facet. The colored points represent the mean activity per hour for each mouse while the dashed and solid curves represent predictions from the naïve model and mixed-model, respectively. The mixed model closely follows the observed data and varies substantively between mice, whereas the naïve model often produces inaccurate predictions. Model fit metrics (AIC, BIC, and RMSE) were lower for the mixed model, indicating superior performance (Table 2).

**Figure 2.**
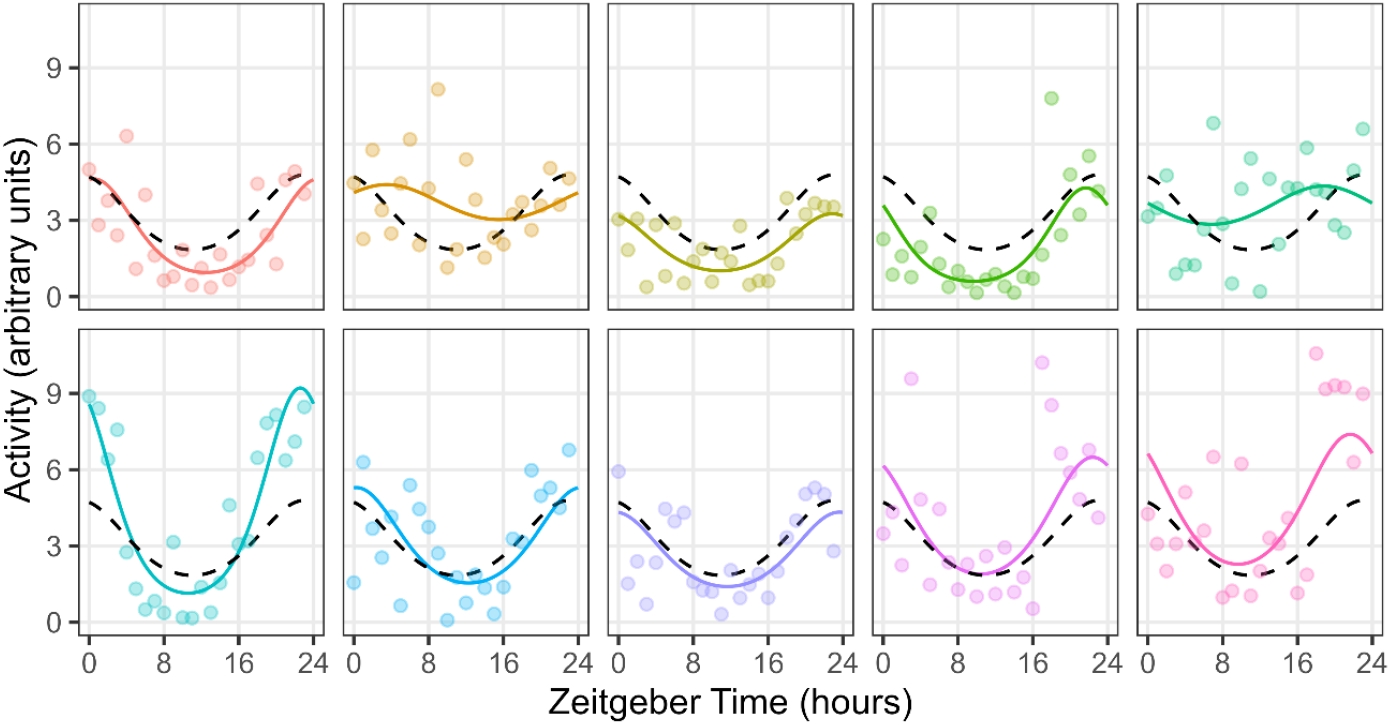
Activity profiles for wildtype (C3H/HeN) mice under constant conditions. Points represent the mean activity during each hour of the day. The dashed line represents the model predictions from the naïve (non-mixed) Poisson model and the colored lines represent the those from the Poisson mixed model.

### Advantages and Applications

GLMMcosinor offers several advantages over traditional cosinor models(9), making it a powerful tool in biostatistics, chronobiology, and clinical research. Its support for diverse data types—including count and zero-inflated data—broadens its applicability. By incorporating mixed-effects modeling, GLMMcosinor improves parameter estimation accuracy by accounting for within-subject correlations.

This flexibility enables applications in gene expression rhythms, hormonal cycles, and other time-dependent biological processes. Additionally, GLMMcosinor provides user-friendly documentation, tutorials, and an interactive Shiny app, making it accessible to researchers at various expertise levels.

### Visualization and Analysis

GLMMcosinor includes intuitive visualization tools that enhance data interpretation. The polar_plot() function effectively compares amplitude and phase shifts across experimental groups by representing amplitude as radius and phase as angle. This visualization is particularly useful in chronobiology, where detecting phase shifts and amplitude differences is crucial for understanding rhythmic variations.

The autoplot() function generates time-series plots that illustrate how rhythms evolve over time. By displaying trends in amplitude, phase, and periodicity, these plots allow researchers to observe rhythmic changes over the study duration. Additionally, users can overlay covariate effects, group responses, and confidence intervals, providing a comprehensive representation of modeled rhythms and experimental factors. This visualization is particularly valuable for assessing subtle differences in treatment effects and ensuring model reliability. Beyond visualization, GLMMcosinor supports diagnostic tools essential for validating model assumptions. Packages such as DHARMa facilitate residual analysis and model validation(22), ensuring that estimated rhythms reflect true biological signals rather than statistical artifacts. By integrating these tools, GLMMcosinor enhances both the interpretability and robustness of cosinor modeling in biological research.

## Conclusion

GLMMcosinor marks a significant advancement in cosinor modeling, offering a flexible, robust, and comprehensive framework for analyzing rhythmic biological data. By integrating Generalized Linear Mixed Models (GLMMs), it overcomes the limitations of traditional cosinor methods, enabling the analysis of non-Gaussian distributions, hierarchical structures, and complex study designs. Its support for diverse data types, hierarchical modeling, and advanced visualization tools makes it a powerful and versatile resource for studying rhythmic processes in biomedical and health sciences.

As a next-generation tool in rhythm analysis, GLMMcosinor has the potential to reshape chronobiology, biostatistics, and clinical research, driving deeper insights into time-dependent biological phenomena. Its user-friendly implementation and extensive documentation make it accessible to researchers at all levels, accelerating discoveries in circadian biology, endocrinology, and beyond.

## Declarations

### Ethics approval and consent to participate

All experimental procedures were approved by The University of Queensland’s Animal Ethics Committee and conducted in accordance with both the Australian code for the care and use of animals for scientific purposes.

### Consent for publication

Not applicable

### Availability of data and materials

The GLMMcosinor code is available on GitHub: https://github.com/ropensci/GLMMcosinor/ and online documentation are accessible here: https://docs.ropensci.org/GLMMcosinor/. GLMMcosinor Shiny App is available via the online documentation site on GitHub. Experimental datasets used and/or analysed during the current study are available from the corresponding author on reasonable request.

### Authors’ contributions

Conceptualization: RP, OR

Methodology: RP, OJ, NW

Resources: RP, OR

Investigation: RP, OJ, OR

Analysis: RP, OJ,

NW Statistics: RP, OJ, NW

Visualization: RP, OJ, OR

Funding acquisition: OR, PC

Writing—original draft: RP, OJ, OR

Writing—review & editing: RP, OJ, NW, PC, OR

## Acknowledgements

Not applicable

